# Contrasting effects of land cover on nesting habitat use and reproductive output for bumble bees

**DOI:** 10.1101/2020.09.26.315044

**Authors:** Genevieve Pugesek, Elizabeth E. Crone

## Abstract

1. Understanding habitat quality is central to understanding the distributions of species on the landscape, as well as to conserving and restoring at-risk species. Although it is well-known that many species require different resources throughout their life cycles, pollinator conservation efforts focus almost exclusively on forage resources.
2. Here, we evaluate nesting habitat for bumble bees by locating nests directly on the landscape. We compared colony density and colony reproductive output for *Bombus impatiens*, the common eastern bumble bee, across three different land cover types (hay fields, meadows, and forests). We also recorded nest site characteristics, e.g., the position of each nest site, for all *Bombus* nests located during surveys to tease apart species-specific patterns of habitat use.
3. We found that *B. impatiens* nests exclusively underground in two natural land cover types, forests and meadows, but not in hay fields. *B. impatiens* nested at similar densities in both in meadows and forests, but colonies in forests had much higher reproductive output.
4. In contrast to *B. impatiens, B. griseocollis* frequently nested on the surface of the ground and was almost always found in meadows. *B. bimaculatis* nests were primarily below ground in meadows. *B. perplexis* nested below ground in all three habitat types, including hay fields.
5. For some bumble bee species in this system, e.g., *B. griseocollis* and *B. bimaculatis*, meadows, the habitat type with abundant forage resources, may be sufficient to maintain them throughout their life cycles. However, *B. impatiens* might benefit from heterogeneous landscapes with forests and meadows. Further research would be needed to evaluate whether hay fields are high-quality nesting sites for the one species that used them, *B. perplexis*.
6. *Synthesis and applications*. In the past, *Bombus* nesting studies have been perceived as prohibitively labor-intensive. This example shows that it is possible to directly measure nesting habitat use and quality for bumble bee species. Applying these methods to more areas, especially areas where at-risk *Bombus* spp. are abundant, is an important next step for identifying bumble bee habitat needs throughout their life cycles.

## Introduction

Ecologists often rely on habitat restoration and conversion projects to aid species recovery or enhance ecosystem services (Long 2009, Tonietto and Larkin 2018). Implementing effective management strategies can be challenging without first identifying the resources and conditions necessary for target populations to persist (Thomas 1980). This general problem is illustrated by initiatives to address declines of pollinating insects. Most efforts to enhance pollinator habitat focus on protecting or enhancing foraging resources, i.e. floral abundance (Decourtye et al. 2010, Dicks et al. 2015). Although these efforts are laudable, increasing forage resources does not guarantee pollinator populations will be maintained. Insect populations may be limited by other factors, including the availably of host plants or nesting material (Potts et al. 2005, Flockhart et al. 2015). At the present time, for many bee species, our understanding of nesting ecology is insufficient to design effective management strategies.

In this study, we evaluate nesting habitat for bumble bees (*Bombus* spp.), a group of economically and ecologically important pollinating insects (Corbet et al. 1991, Carreck and Williams 1998). Knowledge of nesting habitat requirements is a key limiting step in conserving bumble bee populations, despite extensive background data about other aspects of their life cycle, such as foraging (Osborne et al. 1999, Goulson 2010) and physiology (Woodard 2017). For example, in 2020, the US Fish and Wildlife Service decided that it was “not prudent” to designate critical habitat for the endangered rusty-patched bumble bee, in part because of lack of knowledge about its specific nesting and overwintering habitat needs (US Federal Register 2020). In part, this knowledge gap exists because many researchers consider locating *Bombus* nests to be prohibitively difficult (Liczner and Colla 2019). Most studies describing nest locations are anecdotal, describing the location of only a few nests (e.g., Gonzalez et al. 2004). A handful of researchers, mostly in the UK, have used systematic searches to estimate *Bombus* nest densities (e.g., Osborne et al. 2008, O’Connor et al. 2017), but in these studies densities are confounded with search effort. Alternatively, researchers have used indirect methods to study the spatial distribution of *Bombus* nests, i.e. counting nest-searching queens in different habitat types (Svensson et al. 2000, O’Connor et al. 2017) or estimating nest locations from the locations of sibling worker bees (Lepais et al. 2010, Redhead et al. 2015, Carvell et al. 2017). These methods provide some coarse scale data but lack the precision to identify nest locations with the kind of resolution to provide meaningful guidance. For example, past studies have typically found bumble bee nests in a range of habitat types, and have little ability to discriminate use from detectability, or to separate habitat use from habitat quality (cf. Van Horne 1983). Therefore, one interpretation is that bumble bees are generalists with no specific nesting habitat needs (cf. US Federal Register 2020).

In this study, we directly located bumble bee nests in the field and measured variation in reproductive output, to test for differences in both habitat use and habitat quality. Although *Bombus* nests are much harder to find than foraging worker bees, they are not harder to find than some other taxa that have been widely studied with these basic demographic tools, e.g., ground-nesting birds. At a set of landscapes near Ipswich, Massachusetts, USA, we identified nesting habitat use by locating *Bombus* nest sites across three land cover types, using mark-resight methods (Iles et al. 2019) to account for imperfect detection of nests and the possibility of habitat-specific differences in detection probability. After locating nests, we evaluated nesting habitat quality by estimating the reproductive output of colonies of the most common species, *B. impatiens* in different land cover types. Our results demonstrate the feasibility of monitoring *Bombus* nests, and the importance of separating habitat use from habitat quality.

## Materials and methods

### Study sites

This research was conducted at three properties located just outside of Ipswich, Massachusetts, USA, owned and managed by The Trustees of Reservations, a non-profit land trust (Appendix S1). Two of the properties, Appleton Farms (42°38’52.09”N, 70°51’1.01”W) and Appleton Grass Rides (42°38’33.26”N, 70°51’57.12”W), are mixed-use agricultural landscapes. Appleton Farms is an active farm, whose dominant land cover types include hay fields, meadows, forests, and rangeland for cattle grazing. The adjacent site, Appleton Grass Rides, is dominated by forests, though a large meadow and several hay fields are maintained at the property. The third property, Greenwood Farms (42°41’35.77”N, 70°48’59.65”W), is a historic property located approximately 10 km from the other sites that is no longer used for agriculture. This property is dominated by natural areas, including forests, marshes and grassy meadows.

### Nest density surveys

To estimate *Bombus* nest densities, we conducted systematic searches across three land cover types – hay fields, meadows, and deciduous forests. Prior to field work, we noted queen *Bombus* searching for nest sites within meadows and forests, as well as in managed hay fields, and thus limited our study to these land cover types. Hay fields are harvested 2-3 times per year and represent the most intensively managed land cover type. Meadows and forests are not managed intensively, though meadows are mowed annually to prevent succession.

In 2018, we conducted nest density surveys at 30 1500 m^2^ plots. Each land cover type (hay field, meadow, forest) was represented by 10 plots, with 3-4 plots per land cover type located at each property. An additional three plots within hay fields were surveyed at Appleton Farms, as there are no hay fields at Greenwood Farms. To select plots, we first screened each property for potential study sites by excluding areas whose primary vegetation consisted of invasive plants or that were marshy, as bumble bees will avoid nesting in poorly drained areas (Lye et al. 2011). For forested plots, we considered only areas adjacent to hay fields or meadows, and we selected plots within meadows vegetated primarily by grasses, forbs, and some herbaceous shrubs. From acceptable areas, we selected plots haphazardly, unless we had prior knowledge of *Bombus* nest locations, in which case plots were chosen randomly. In 2019, we selected 30 different plots in the same manner, for a total of 60 survey plots. We searched each plot for an hour-long period once a week, for a total of four sampling occasions per plot. To locate nests, a single researcher (G. Pugesek) walked slowly through each plot, searching for worker traffic around nest entrances. Once a potential nest site was located, we confirmed the presence of a nest by waiting for at least 4 workers to exit or enter (Rao and Skyrm 2013). The first time a nest was located, the nest entrance was marked with an inconspicuous, numbered metal plant tag, and a single worker from each nest was collected to identify the colony to species (Iles et al. 2019). During subsequent searches, we recorded if the nest was re-sighted to generate a capture history for each nest. Nest searches were carried out when *B. impatiens* colonies were large and when worker traffic at nest entrances was noticeable: from July 13^th^ to August 15^th^ in 2018 and July 12^th^ to August 14^th^ in 2019 (Iles et al. 2019). All surveys were conducted between 8:00 AM and 6:00 PM when the weather was clear.

### Free searches for bumble bee nests

To maximize the number of nests located to make comparisons of habitat use across species, we supplemented our nest density surveys by conducting “free searches” for bumble bee nests, adapted from the methodology described by O’Connor et al. (2012). Searches were conducted by four different investigators in natural areas, e.g., dry meadows, field margins, and forests, at Appleton Farms and Grass Rides in 2018 and at all three properties in 2019. Free searches for bumble bee nests took place prior to nest density surveys, between mid-April and mid-July, because the colonies of *B. griseocollis* and *B. perplexus* began to senesce prior to our last nest density surveys. Some of the same areas were searched during nest density surveys and free searches for nest sites; however, there was not much evidence that the probability we would locate nests during nest density surveys was influenced by prior knowledge of nest sites (see Appendix S2).

While conducing free searches for nests, researchers walked haphazardly through search areas at their own pace, searching for worker traffic around nest site entrances. After we located a potential nest site, we confirmed the presence of a nest by waiting for the queen bumble bee to return with pollen baskets, or for at least 2 workers to either exit or enter the nest. The length of searches depended on weather conditions, but generally ranged from 2-3 hours. In 2018, we did not record the time spent searching each land cover type, though, in general, we spent more time searching for nests in open areas. In 2019, we spent approximately 150 hours searching for nests in meadows and 90 hours searching for nests in forests. All surveys were conducted between 8:00 AM and 6:00 PM when the weather was clear.

### Assessing nest site positions

The position of nest sites, e.g., whether the nest was located on the ground surface or below ground, was confirmed by visually inspecting nest site entrances, excavating nests, or by gently tapping the vegetation around nest sites (Kupchikova 1960). For most subterranean nests, the nest was clearly located underground, though removing some leaf material or vegetation was sometimes necessary to expose the entrance. For nests on the surface of the ground, in 2018, we conducted a similar visual inspection of the nest sites, i.e., we recorded whether worker bees seemed to be entering a tussock of grass or a rodent nest on the surface of the ground. In 2019, we excavated all surface nests after the colony had expired, or gently tapped the surface of the ground around the nest site with a stick, listening for buzzing workers (Kupchikova 1960).

### Monitoring gyne output for each nest

We monitored *B. impatiens* nest entrances for activity of newly emerged gynes (i.e., female social insects with the potential to become queens) by watching nest entrances for 30 minutes, 2 times a week (see Appendix S3 for discussion of this protocol compared to others). We used colonies we had encountered during nest density surveys and free searches for nests, as described above, as well as three colonies encountered incidentally during field work to maximize our sample size. We ensured colonies had not expired prior to reproduction by monitoring each colony for activity (at least one worker either entered or exited the colony entrance during a one-hour period) prior to the surveys. Nest entrances were monitored from August 27^th^ to October 23^rd^ in 2018. No gynes were observed at field sites prior to this date. In 2019, we watched nest entrances from August 5^th^ to October 30^th^, as we began observing gynes at nests in early August.

During each 30-minute survey period, gynes entering or exiting the nest were netted, cold-anesthetized, and marked with a unique number tag (Queen marking kit, BetterBee). At the end of each sampling period, captured gynes were released. Each colony was monitored until no workers or reproductives were encountered for 4 subsequent observation periods. Surveys took place between 9:30 AM and 5:00 PM; the time of day each nest was sampled was staggered. To quantify reproductive output for each nest, we tallied the total number of unique gynes encountered over the entire study period at each nest. We encountered no gynes at several nest sites; thus, using mark-recapture methods to estimate the total number of gynes produced by these colonies was not feasible. However, for colonies that produced gynes, we confirmed that the observed number of unique gynes encountered at each nest site was a strong positive predictor of the number of gynes produced by each colony as estimated using mark-recapture methods (*R*^2^ = 0.946, *P* < 0.001, see Appendix S4).

### Statistical methods

*B. impatiens* nest densities were estimated using mark-resight methods. We used closed population models to estimate capture probabilities and nest densities from *B. impatiens* capture histories, because bumble bee nest site locations are fixed, and few colonies expired during our surveys. We implemented a Bayesian analysis using spatially stratified closed population models (Kéry and Royle 2015). Choosing a Bayesian analysis allowed us to estimate detection probability and account for the variation in abundance between survey plots when estimating nest densities, a model structure that is not available in standard maximum likelihood mark-recapture software, e.g., program MARK (White and Burnham 1999). Models were run in JAGS using the R package jagsUI (Kellner 2019) for 300,000 interactions across 3 chains with a burn-in period of 10,000 iterations. We fit models including differences in detection among years and landcover type and performed model selection by estimating posterior model weights using indicator variables (Kuo and Mallick 1988, Kéry and Royle 2015). Estimates of posterior model weights are sensitive to model priors; thus, we set priors to posterior distributions calculated using the full model (Aitkin 1991, Kéry and Royle 2015). We verified models had converged by inspecting trace plots visually and confirming Gelman-Rubin convergence diagnostic values were less than 1.05 (Gelman and Rubin 1992, Kéry and Royle 2015). To determine if estimates of nest density differed across land cover types in each year of the study, we calculated a posterior distribution of the difference between means. If the 95% credible intervals of the difference overlapped with zero, we considered the means to differ. As *B. impatiens* nests were not found in hay fields, we assumed the same detection probability for nests in hay fields and meadows when estimating nest densities. In the absence of any nests in hay fields, it is impossible to statistically estimate capture probabilities. However, it is reasonable to expect that our ability to detect nests would be similar between hay fields and meadows, because both land cover types are dominated by grasses and herbaceous plants.

We assessed species-specific patterns of habitat use using multinomial logistic regression models, fit using the multinom function in the R package nnet (Venables and Ripley 2002; this and other R analyses were implemented in R version 3.6.1 (R Development Core Team 2019).). We pooled data from across years and study sites, and assessed nests located during free searches and nest density surveys as well as nest sites found incidentally during other field work. Only nests that produced workers were included in this analysis, as we were unable to reliably identify queen bumble bees to species via observations in the field. We used separate models to compare nest position and surrounding land cover across bumble bee species. Species, our predictor variable, was categorized into 3 groups: *B. impatiens, B. griseocollis*, and other. *Bombus* species categorized as “other” (*B. perplexus, B. bimaculatus*, and *B. vagans*) were not encountered frequently enough to be assessed independently. Likelihood ratio tests, implemented using the command lrtest in package lmtest (Zeileis and Hothorn 2002), were used to compare univariate and null models. Post-hoc comparisons of estimated marginal means were made using the R package emmeans.

We analyzed differences in reproductive output, i.e., the total number of unique gynes encountered at each nest site, for *B. impatiens* using generalized linear models with negative binomial error distributions to account for overdispersion. Data were pooled across study sites due to low sample sizes. Models were fit using the R package MASS (Venables and Ripley 2002). Fixed effects included the surrounding land cover type (forest or meadow) and the year the colony was monitored (2018 or 2019); we also included an interaction between land cover and year. Candidate models included all possible combinations of variables and interactions (Table 1). Models were ranked and compared using Akaike’s information criteria (AIC_*C*_) corrected for small sample size using the AICc function in R.

**Table 1.** AICc statistics for competing generalized linear models (family = negative binomial) used to estimate reproductive output of *B. impatiens* colonies.

| Model abbr. | Effects | df | AICc | Model weight |
| --- | --- | --- | --- | --- |
| $M_0$ | *Null* | 2 | 200.6 | 0.203 |
| $M_c$ | *Cover* | 3 | 200.0 | 0.264 |
| $M_y$ | *Year* | 3 | 201.7 | 0.116 |
| $M_{c+y}$ | *Cover + Year* | 4 | 199.6 | 0.332 |
| $M_{c*y}$ | *Cover\*Year* | 5 | 202.3 | 0.085 |

## Results

### Nest densities

In two years, we located 29 *B. impatiens* nests during nest density surveys. *B. impatiens* nests were not found in hay fields, while in meadows and forests, we found 17 and 12 *B. impatiens* nests, respectively. We located bumble bee nests of other species as well (17 *B. griseocollis*, 1 *B. bimaculatus*, 2 *B. vagans*, and 2 *B. perplexus* nests), for a total of 51 nests (Appendix S5: Table S1).

We estimated capture probabilities for *B. impatiens* nests in meadows and forests, as no *B. impatiens* were found in hay fields. The model with the highest posterior model weight included land cover type as a covariate, *M*_*c*_, with a posterior probability of 0.519. Detection probability was lower in forests (0.335) than in meadows (0.550). In contrast, the null model had a low posterior probability (0.174). There was not much evidence our ability to locate nests differed between survey years, as the models which included year, *M*_*c*_, and both land cover type and year, *M*_*c+y*_, as covariates had low posterior probabilities (0.098 and 0.296, respectively). This covariate was thus excluded from further analyses.

In 2018 and 2019, estimated nest densities in hay fields were lower than in meadows and forests (Fig. 1). In 2018, nest densities were lower in meadows than forests, with estimated nest densities of 4.28 nests · ha^-1^ in meadows and 7.58 nests · ha^-1^ in forests (a difference of −3.29 nests · ha^-1^; 95% credible intervals [CRI]: −10.50 - 2.45). In 2019, we saw a larger difference in the opposite direction (+4.67 nests · ha^-1^, 95% CRI: −0.61 - 10.23), with estimated nest densities of 7.41 nests · ha^-1^ in meadows and 2.74 nests · ha^-1^ in forests. Nest densities in the meadow were lower in 2018 than 2019 (−3.13 nests · ha^-1^ 95% CRI: −8.80 - 2.16), whereas nest densities in the forest were higher in 2018 than 2019 (4.84 nests · ha^-1^, 95% CRI: −0.60 - 11.61).

**Fig. 1.**
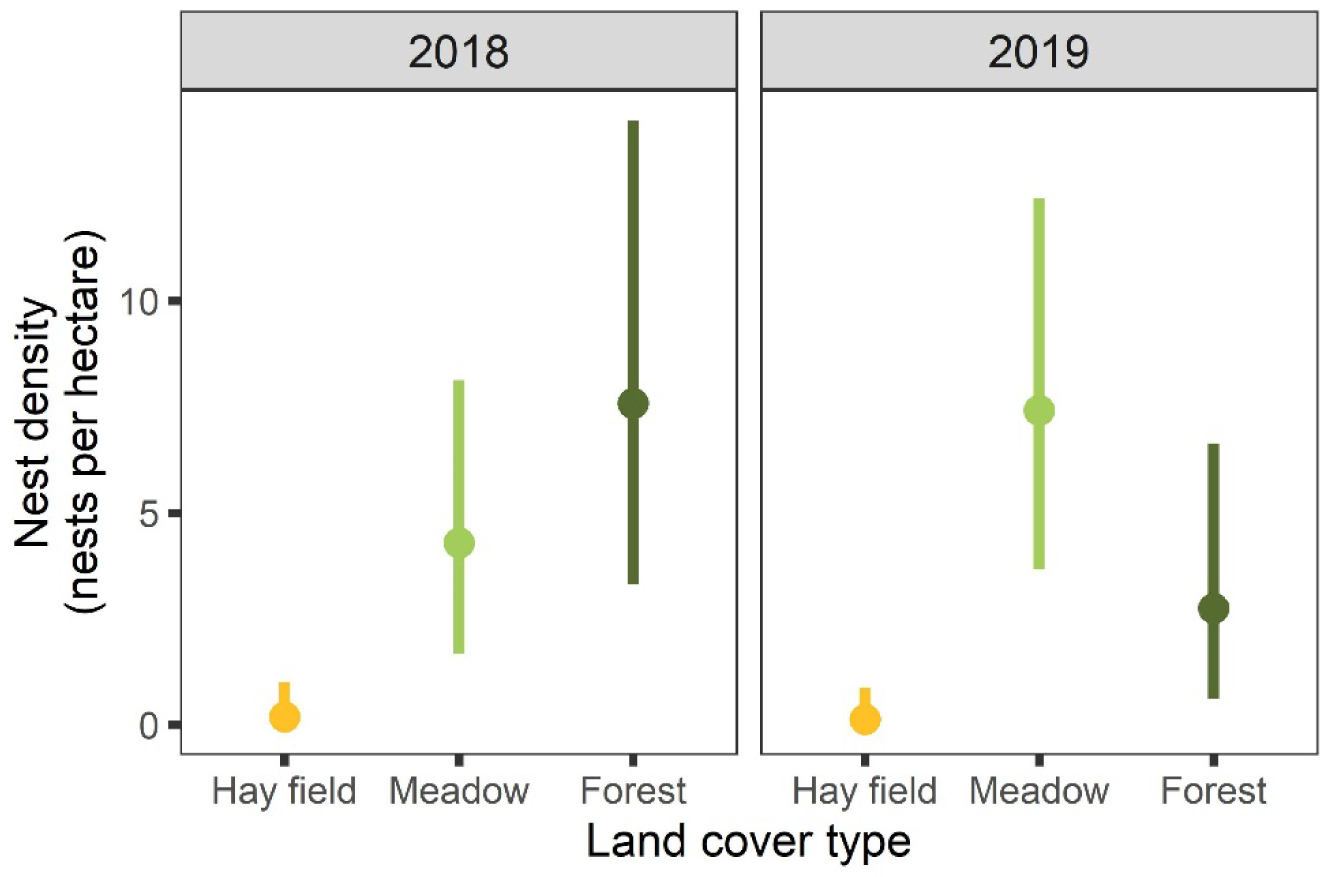
Nest densities and associated 95% Bayesian credible intervals for *B. impatiens* across three different land cover types estimated using spatially explicit closed capture models.

### Species-specific patterns of habitat use

A total of 122 bumble bee nests were located via a combination of free searches, nest density surveys, and incidental encounters. Most bumble bee nests located were subterranean (61.9%) though many nests were also found on the surface of the ground, under tussocks of grass or within abandoned rodent nests (36.4%). A handful of nests were found beneath decaying stumps. Most nests sites were found in meadows (83.5%) as opposed to forests (14.8%) or hay fields (1.6%) (Fig. 2), which was expected, given greater search effort in open natural areas during free searches.

**Fig. 2.**
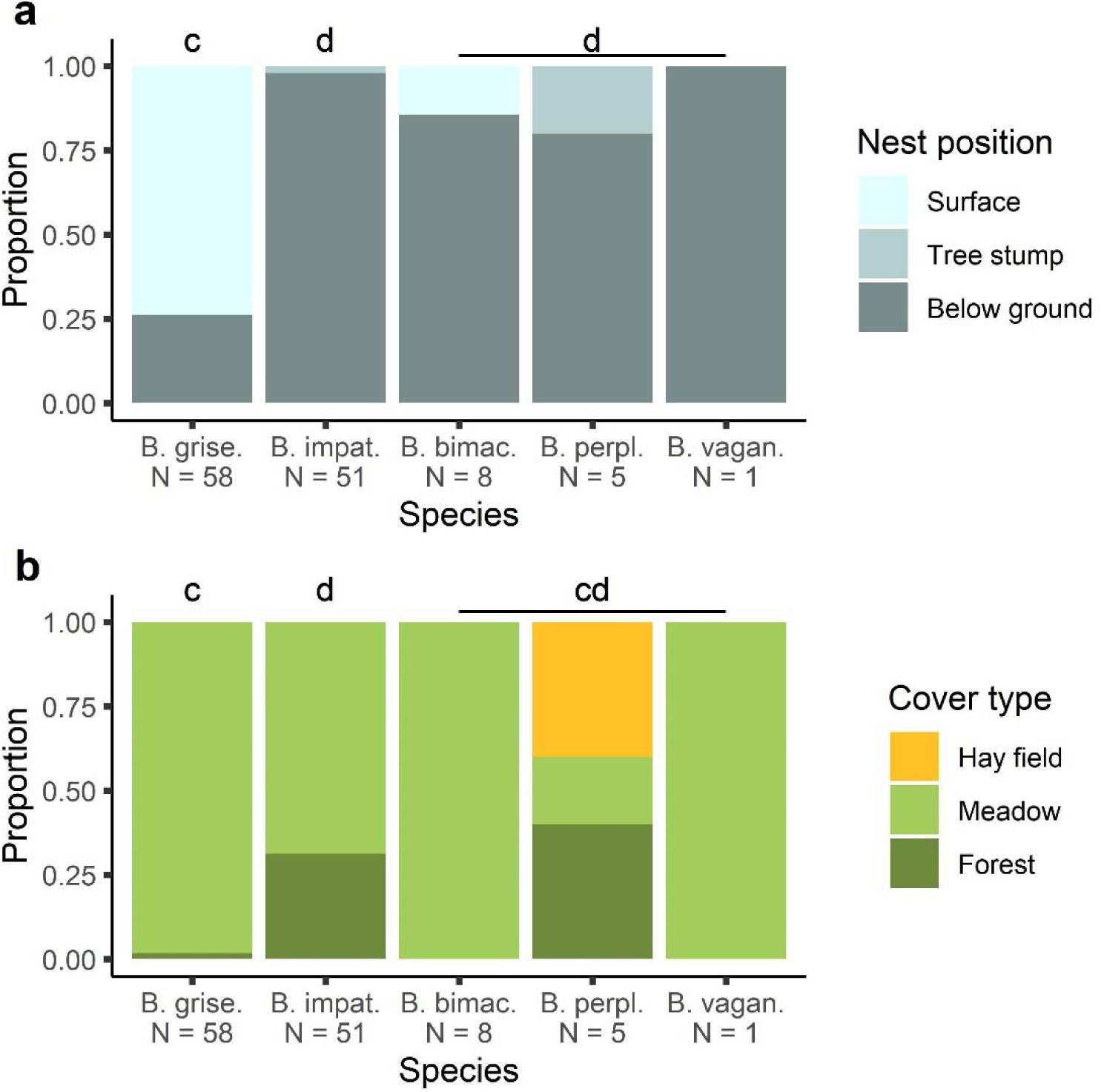
a) The position of nest sites and b) the land cover type where nests were found for *B. griseocollis* (B. grise.), *B. impatiens* (B. impat.), *B. bimaculatus* (B. bimac.), *B. perplexus* (B. perpl.), and *B. vagans* (B. vagan.). Letters (c and d) indicate statistical significance groups within each subpanel.

*Bombus* species differed in terms of nest site position (*χ*^2^ = 85.69, df = 4, P < 0.001). Compared to *B. impatiens* and the other *Bombus* spp., a greater proportion of *B. griseocollis* nested on the surface of the ground (73.7%), while similar proportions of *B. impatiens* nests (0.0%) and the nests of other *Bombus* spp. (7.0%) were found on the ground surface (Appendix S6: Table S1). A greater proportion of *B. impatiens* and the other *Bombus* spp. nested below ground compared to *B. griseocollis* (26.3%), though similar proportions of *B. impatiens* (98.0%) and other *Bombus* spp. (84.6%) nested below ground (Appendix S6: Table S1). The proportion of nests found under tree stumps was similar for *B. impatiens* (2.0%), *B. griseocollis* (0.0%), and other *Bombus* spp. (7.7%, Appendix S6: Table S1).

The use of different land cover types while nesting also differed across *Bombus* species (*χ*^2^ = 29.79, df = 4, P < 0.001). Relative to *B. impatiens*, for which 68.6% of nests were found in meadows and 31.4% of nests were found in forests, a greater proportion of *B. griseocollis* nests were found in meadows (98.3%) and a lesser proportion were found in forests (1.7%, Table Appendix S6: Table S2). No other comparisons were statistically significant (Appendix S6: Table S2).

### Reproductive output

In 2018, we monitored 8 *B. impatiens* colonies nesting in meadows and 9 *B. impatiens* colonies nesting in forests, and in 2019 we monitored 13 *B. impatiens* colonies nesting in meadows and 4 *B. impatiens* colonies nesting in forests. We collected and marked a total of 271 unique gynes from nest entrances over the course of two years.

Colony reproductive output was higher in forests than in meadows, and colonies reproduced more in 2019 than 2018 (Fig. 3). The highest-ranking model included additive effects of both land cover and year (Table 1), though the model including only an effect of land cover (dAICc = 0.5) and the null model also ranked highly (dAICc = 1.0, for all other model comparisons, dAICc > 2). Colonies found in forests produced nearly three times as many gynes (14.0 gynes encountered · nest^-1^) as colonies found in meadows (5.0 gynes encountered · nest^-1^). Colonies also produced about twice as many gynes in 2019 (10.9 gynes encountered · nest^-1^) than in 2018 (5.5 gynes encountered · nest^-1^).

**Fig. 3.**
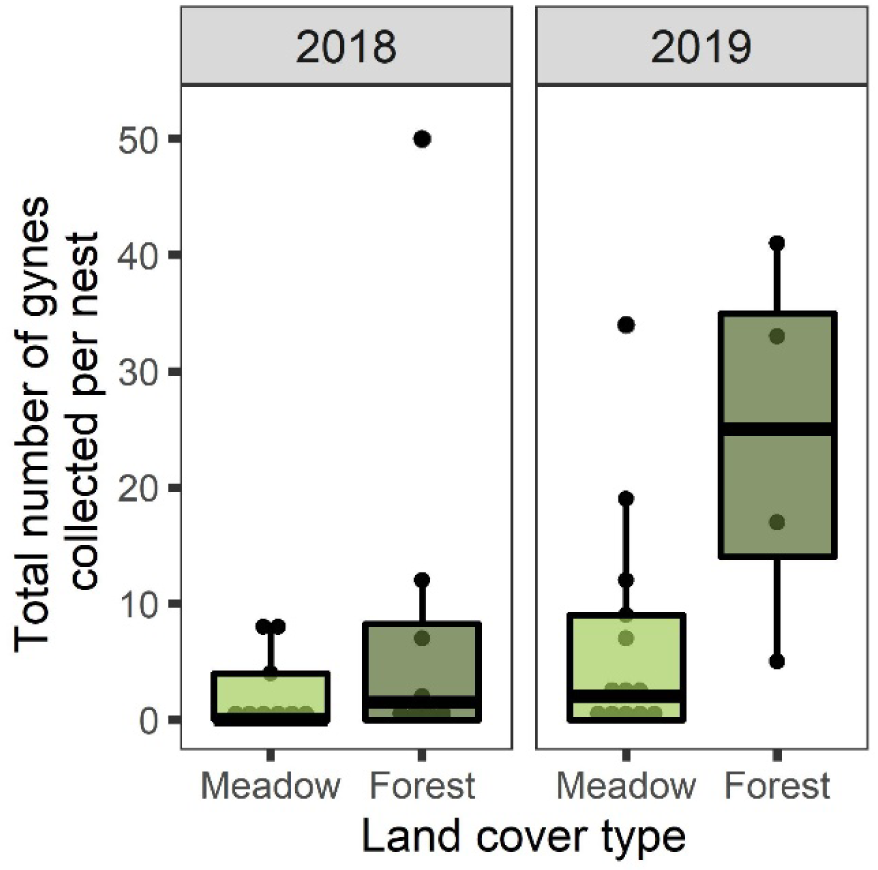
The total number of unique gynes encountered at *B. impatiens* colonies located in forests and meadows.

## Discussion

Forests and meadows provided the conditions necessary for *B. impatiens* to nest: in fact, over the course of our nest density surveys, we found a similar number of *B. impatiens* nests in forests (N = 12) and in meadows (N = 17). However, *B. impatiens* colonies found in forests produced more gynes than those found in meadows, suggesting forests are higher quality nesting habitat for *B. impatiens* relative to meadows. We found no *B. impatiens* nests in hay fields, though queens were observed searching for nests within this land cover type prior to surveys. Human activity, such as tilling or mowing, may destroy potential nest sites or drive away the animals whose burrows would be later used by nesting bumble bees. Bumble bee nests are typically found in natural or semi-natural areas, e.g. hedgerows adjacent to croplands, as opposed to highly modified areas (Liczner and Colla 2019). Worker bumble bees are also abundant and are more likely to visit flowers in crop lands adjacent to natural habitats, suggesting bumble bees use these more natural areas to nest (Greenleaf and Kremen 2006, Morandin et al. 2007).

In some contexts, there is a tendency to treat “bumble bees” as a single taxon (Becher et al. 2018), even though it is a genus with a range of life history variation among species. Our study shows that areas that provide suitable nest sites for *B. impatiens* may fail to provide nest sites for other species. While *B. impatiens* nest above ground with similar nest densities in forests and meadows, *B. griseocollis* nests were almost exclusively on the surface in meadows (see Fig. 2 and Appendix S5). Other *Bombus* species, e.g. *B. vagans* and *B. bimaculatus*, were also encountered in open habitats (Fig. 2). Past studies have noted species-specific land use, e. g. some species of bumble bees tend to use nest sites on the ground surface, while other bee species tend to nest below ground (Lye et al. 2012). Other authors have noted, albeit anecdotally, *B. impatiens* nesting below ground (Plath 1922) and *B. griseocollis* nesting on the ground surface (Harder 1986). However, our study is the first to systematically compare nesting habitat of North American *Bombus* (i.e., to survey nests in a way that is amenable to making statistical comparisons, without relying on the use of artificial nest boxes).

We found that detection probability was higher in meadows than forests in our study system. Therefore, we might have underestimated the importance of forests if we had searched study plots for nests only once. Several past studies have compared nest densities between forests and grasslands without correcting for nest detection probabilities (Osborne et al. 2008, O’Connor et al. 2017). However, even very small differences in detection probability across treatments or environmental gradients can lead to incorrect conclusions if unaccounted for (Archaux et al. 2012). Many recent studies of bumble bee nesting rely on the effort of citizen scientists (Osborne et al. 2008, Lye et al. 2012) or trained dogs (Waters et al. 2011, O’Connor et al. 2012) to locate bumble bee nests, and thus it would be valuable to evaluate whether other survey methods are robust to bias created by imperfect detection.

We observed higher reproductive success of colonies in forests compared to meadows, emphasizing the longstanding notion that habitat use is not always correlated with habitat quality (Van Horne 1983). The question remains as to why *B. impatiens* nesting in forests were found to produce more queens than those nesting in meadows. *B. impatiens* may face more competition for preferred nest sites in meadows as opposed to forests, as we encountered *B. griseocollis* more frequently in meadows (Table 4). If bumble bees are nest site limited (McFrederick and LeBuhn 2006, Inoue et al. 2008), competition may drive *B. impatiens* queens to accept lower quality nest sites to establish colonies at all. Abiotic conditions in forests may have been more suitable for nesting *B. impatiens*, as tree cover can buffer air and soil temperatures and offer protection from the elements (Chen et al. 1995, Gaudio et al. 2017). Another possibility is that detection probability depended on colony size. Because detection probability is higher in meadows, we may have been able to locate more small colonies. Bumble bee queen production increases with colony size (Crone and Williams 2016, Goulson et al. 2018) and therefore a detection bias against small colonies in forests could lead to estimated lower average queen production in meadows. Associating detection probability with colony size could be an interesting avenue for future research.

Reproductive success of colonies varied across years as well as among habitat types (Fig. 3). Temporal variation in reproductive success has been previously observed for bumble bees: Richards (1978) found 57% of bumble bee colonies nesting in artificial domiciles in to produce sexuals in 1970, while only 16% of colonies produced sexuals the following year. Similarly, a study monitoring the reproductive output of wild bumble bee colonies in the UK found that 71% of colonies produced gynes in 2010, while only 21.1% of colonies produced gynes in 2011 (Goulson et al. 2018). While the cause of this variation is not always clear (Goulson et al. 2018), bumble bee population dynamics are likely impacted by factors such as precipitation and temperature (Goodwin 1995).

Although we observed higher reproductive output of *B. impatiens* nests in forests, the forested habitat patches in our study system were highly fragmented. Nests were close to meadow edges (often less than 10 m) compared to bumble bee forage distances (routinely over 1.5 km, see Osborne et al. 2008). Forests provide few floral resources in our study region, but almost all study plots in forests were adjacent to areas where floral resources were abundant, e.g. wet meadows (Appendix S1). Bumble bee nest densities are higher at forest edges than in core forest in the UK (Osborne et al. 2008), and nest searching bumble bee queens are less abundant in forests than at forest edges (Svensson et al. 2000). Vaudo et al. (2018) also suggested bumble bee colonies placed in core forest forage less and are less successful than colonies placed at forest edges. Therefore, we hypothesize that although forests are higher quality nesting habitat, bumble bees need open habitat for foraging in order to persist at landscape scales. In other words, the important advance of the present study over our previous understanding of *B. impatiens* habitat needs is that forests contribute to population viability by providing nesting habitat, even though they do not contain many floral resources. Both habitat types in proximity are likely necessary for maintaining viable populations.

For many species of bumble bees – especially those that are rare or threatened – habitat requirements during crucial life history events like nesting are poorly understood. While the results presented here primarily represent two common bumble bee species, our findings inform future efforts to monitor bumble bee populations by showcasing feasible strategies for collecting demographic data. Even for these two species, our research emphasizes that bumble bee nesting habitat preferences are species specific: *B. griseocollis* would be more likely than *B. impatiens* to persist in landscapes that contained only meadows, but, because it is surface nesting, would be more susceptible to disturbances like fire. Applying these methods to other landscapes, perhaps especially in regions where at-risk species are still locally common, is a crucial next step in bumble bee conservation.

## Authors’ Contributions

G.P. led field work, while both G. P. and E. E. C., conceived ideas and designed methodology, conducted data analyses, and contributed to writing this manuscript. Both authors approve the final version of this manuscript for submission.

## Acknowledgements

Funding for this research was provided by the Tufts University College of Arts and Sciences and the National Science Foundation (award DEB 1354022 to E. Crone). We would like to acknowledge Maria Ostapovich, Madeline Bondy, and Stuart Farnham for their assistance in the field. We thank James Michielini for his assistance converting raw data into capture histories for gynes and completing the analyses described in Appendix S4, and Dr. Andy Royle for his assistance implementing spatially stratified closed population models. We thank The Trustees and Russell Hopping for their support. The authors declare no conflict of interest.

## Notes

### Competing Interest Statement

The authors have declared no competing interest.

### Summary of Updates

Reference added and typo fixed on page 17

